# Ecological risk-taking across age in Barbary macaques

**DOI:** 10.1101/2024.07.19.603766

**Authors:** Tiffany Claire Bosshard, Roger Mundry, Julia Fischer

## Abstract

When foraging, animals face decisions regarding unpredictable resources and varying predation risk, in which the cost-benefit trade-off must be assessed. Life-history theory focuses on resource allocation with regard to development and suggests there should be a variation in risk-taking propensity across age, manifested by a shift in goals from gain acquisition to loss aversion over time. Consequently, we aimed to assess age-related variation in decision-making strategies under risk in Barbary macaques (*Macaca sylvanus*) living at “La Forêt des Singes” in Rocamadour, France. We experimentally varied the perceived risk and potential gains in a 2×2 design. The high-risk stimulus was a rubber snake; the low-risk stimulus a cube with a snakeskin pattern. The high-value food reward was a peanut; the low-value food reward a popcorn. In 2022, we presented 83 subjects with one of the four possible factor combinations, and in 2023, we presented 70 individuals with the inverted combination (e.g. snake-popcorn in 2022, cube-peanut in 2023). We took each monkey’s propensity to retrieve the reward as a measure of risk-taking, and the latency to retrieve the reward as motivation to engage in the risky choice. Regardless of age, subjects generally retrieved the reward when placed next to the cube, whether peanut or popcorn, but were less likely to retrieve the reward placed next to the snake when this was popcorn rather than peanut. However, older individuals were faster than younger conspecifics to retrieve the reward when it was popcorn, whether paired with the cube or snake. Our results do not comply with the notion that risk-taking propensity declines with age, but rather suggest that in this paradigm, the animals assess the costs and benefits of a risky situation across age, and that the subjective value of a resource may change with age.

## INTRODUCTION

In each decision, the cost-benefit trade-off must be assessed. In this sense, decisions may involve a certain level of risk due to the uncertain payoff that an investment might incur (Yates, 1992). While an overly risk-prone strategy may result in losses, injury, or fatality, an overly conservative strategy may result in a missed opportunity to acquire resources (Kacelnik & Bateson, 1996). Risk-taking propensity may vary according to the type and quantity of resources at stake, but has also been shown to vary across populations, groups, and with individual attributes. In humans for instance, cross-national differences in risk-taking were reported in various surveys assessing participants’ willingness to wager monetary stakes (e.g., Rieger et al., 2015; Vieider et al., 2012). Namely, participants from wealthier countries tend to be more risk averse than participants from poorer ones. Gender differences in risk-taking were found as well, with men being more prone than women to initiate conflicts (Campbell, 1999), more likely to engage in activities with a possibility of physical harm (Byrnes et al., 1999), and more willing to take financial risks (Charness & Gneezy, 2012). There are also individual traits that have been shown to influence risk-taking tendencies such as impulsivity, sensation-seeking, and low self-control which are associated with more risky behaviour (Mishra & Lalumière, 2011).

From an evolutionary perspective, life-history theory provides a framework for explaining variation in risk-taking tendencies. Life-history theory focuses on how organisms allocate their resources with regard to the trade-offs that they face during their development, which can influence their behaviour and decision-making strategies (Stearns, 1992). Life-history theory predicts variation in risk-taking propensity over time, with a shift in goals from gain acquisition to loss aversion with age (Machluf & Bjorklund, 2015). Risk-taking at a young age is considered to be adaptive because there is much to gain and little to lose. Conversely, individuals that have survived into old age have little left to gain and have a future that is more predictable, and they would therefore avoid taking risks (see also Gardner & Steinberg, 2005). In humans, it is well established that there is a decline in propensity for risk across the lifespan (Defoe et al., 2015; Willoughby et al., 2021). More specifically, propensity for risk increases in adolescence, peaks in young adulthood and then declines with age. This is true across multiple domains since, as a general rule, adolescents and young adults are more likely than adults over 25 to engage in sensation-seeking activities, substance abuse such as binge drinking, reckless driving thus having more fatal or serious automobile accidents, or take up violent or criminal behaviour (reviewed in Steinberg, 2008). A decrease in risk-taking then manifests across adulthood, with older age groups showing yet a steeper decline in both financial and health risk-taking propensity, and a cessation to engage in risky recreational activities (Josef et al., 2016).

While in humans risk-taking is often considered in the context of perilous activities or economic behaviour, other animals must also make assessments of value and cost-benefit trade-offs regarding resources such as food or mates (Dammhahn & Almeling, 2012). Animals face situations such as foraging for unpredictable resources and navigating environments with varying predation risk, in which erroneous decisions can have deleterious consequences. As such, considering the biological relevance of decision-making under risk, risk-taking tendencies are being studied in animals as well, most prominently in nonhuman primates (hereafter primates). Much like the findings described in humans, risk-taking propensity in primates tends to vary with various factors including species, sex, or individuals’ internal state (reviewed in De Petrillo & Rosati, 2021). For example, bonobos (*Pan paniscus*) are reportedly more averse to risk than chimpanzees (*Pan troglodytes*) in food gambling tasks (Heilbronner et al., 2008; Rosati & Hare, 2011), which was suggested to be attributed to the two species’ varying feeding ecologies. In terms of sex differences, male chimpanzees were described as being more risk-seeking than females across a series of risk domains, as assessed both through observer ratings as well as behavioural choice experiments (Haux et al., 2023). Risk-taking may also be modulated by state-dependent choices, as demonstrated in rhesus macaques (*Macaca mulatta*), becoming more risk-seeking when their energy level is higher (Yamada et al., 2013).

In terms of age-related variation in risk-taking in primates, recent findings suggest that chimpanzees in their adolescent and young adulthood years are more willing to take risks as compared to their adult conspecifics in tasks involving variability in payoffs (Haux et al., 2023; Rosati et al., 2023). Other studies investigating the same concept were less informative in this regard due to small sample sizes with limited age ranges (e.g., tufted capuchins *Sapajus spp.*, De Petrillo et al., 2015; bonobos, Heilbronner et al., 2008; red-capped mangabeys *Cercocebus torquatus torquatus*, Rivière et al., 2018). Therefore, further studies on larger age-heterogeneous populations of primates are essential to pinpoint possible patterns of age-related changes in risk-taking strategies in primates.

In the current study, we aimed to assess risk-taking across age in Barbary macaques (*Macaca sylvanus*) as tested by means of an ecological foraging task. Following life-history theory, we predicted that monkeys would display a higher likelihood to take ecological risks in adolescence and young adulthood compared to mid-adulthood, which would thereafter be followed by a gradual decline in risk-taking propensity over time. We further predicted that the time to decision (latency) would be longer in older individuals than younger ones, reflecting lengthier deliberation times to engage in a risky choice (Deakin et al., 2004). Given the differences in life-history between the sexes (Tarka et al., 2018) and the risk attitudes associated with male dispersal in this species, we also predicted that males would be more willing to take risks than females.

## METHODS

### Study Site and Subjects

We conducted this study on the population of Barbary macaques living at “La Forêt des Singes” in Rocamadour, France. The enclosure is a 20-ha park in which the monkeys range freely and live outdoors year round. In addition to feeding on the natural vegetation, they are provisioned with fruits, vegetables, and cereal every day, and water is available *ad libitum*. The park is open to visitors, who are able to observe the animals from designated walking paths. The population consisted of 146 monkeys in 2022 to 147 monkeys in 2023, each year corresponding to one of two study periods. The animals lived in three naturally formed social groups of about 50 individuals each. The monkeys are habituated to behavioural observations by researchers, and all individuals are recognisable by their inner-thigh tattoo and/or by distinctive physical traits such as individual facial skin pigmentation (de Turckheim & Merz, 1984).

In view of the availability of food, veterinary care if needed, and the lack of predation, the animals are able to grow very old, which allows for a study population that comprises a large age range. Following earlier studies on the same population (Rathke & Fischer, 2021; Rathke et al., 2022), monkeys were considered ‘adolescents’ and ‘young adults’ up to the age of 10 years, ‘middle-aged’ between 11 and 19 years, and ‘old’ from 20 years onwards. Taken together, 88 individuals (48 females, 40 males) from two of the social groups, ranging in age from 3 to 31 years, participated in one or both study periods.

### Procedure

We conducted a behavioural experiment in which we presented pairs of stimuli to the monkeys that varied in terms of their potential risks and gains. We used a two-by-two factorial design involving high- and low-risk stimuli, and high- and low-quality food rewards (Figure 1). The high-risk stimulus was a small rubber snake; the low-risk stimulus was a cube with the same pattern as the snake (see Videos 1 to 5). The high-quality food reward was a peanut; the low-quality food reward a piece of popcorn. Our choice for the snake as high-risk stimulus was based upon the knowledge that snakes, as prominent predators of mammals evolutionarily speaking, have exerted pressure on primates to develop a particularly important anti-predatory response towards them (Isbell, 2006). Regarding the low-risk stimulus, we designed the cube with a snakeskin pattern in order to present the monkeys with a more abstract shape, while following evidence that the scales alone constitute a certain aversion for primates (Isbell & Etting, 2017). In terms of reward quality, we conducted food preference tests on the third, and otherwise untested, monkey group in the park, and found a group-level preference for peanut over popcorn. The goal was to assess whether the animals would retrieve a food reward placed next to a risky stimulus depending on 1) the level of risk of the stimulus and 2) the quality of the reward, and whether the risk-taking would vary across age. The stimuli were presented to the animals in a wooden box (L: 60cm, W: 30cm, H: 40cm). The box was open on one side; this open side was initially covered with a curtain that was held up with a string by the experimenter. During the experiment, the curtain would be lowered for each subject, allowing them to view and have access to the stimuli and participate in the task.

**Figure 1.**
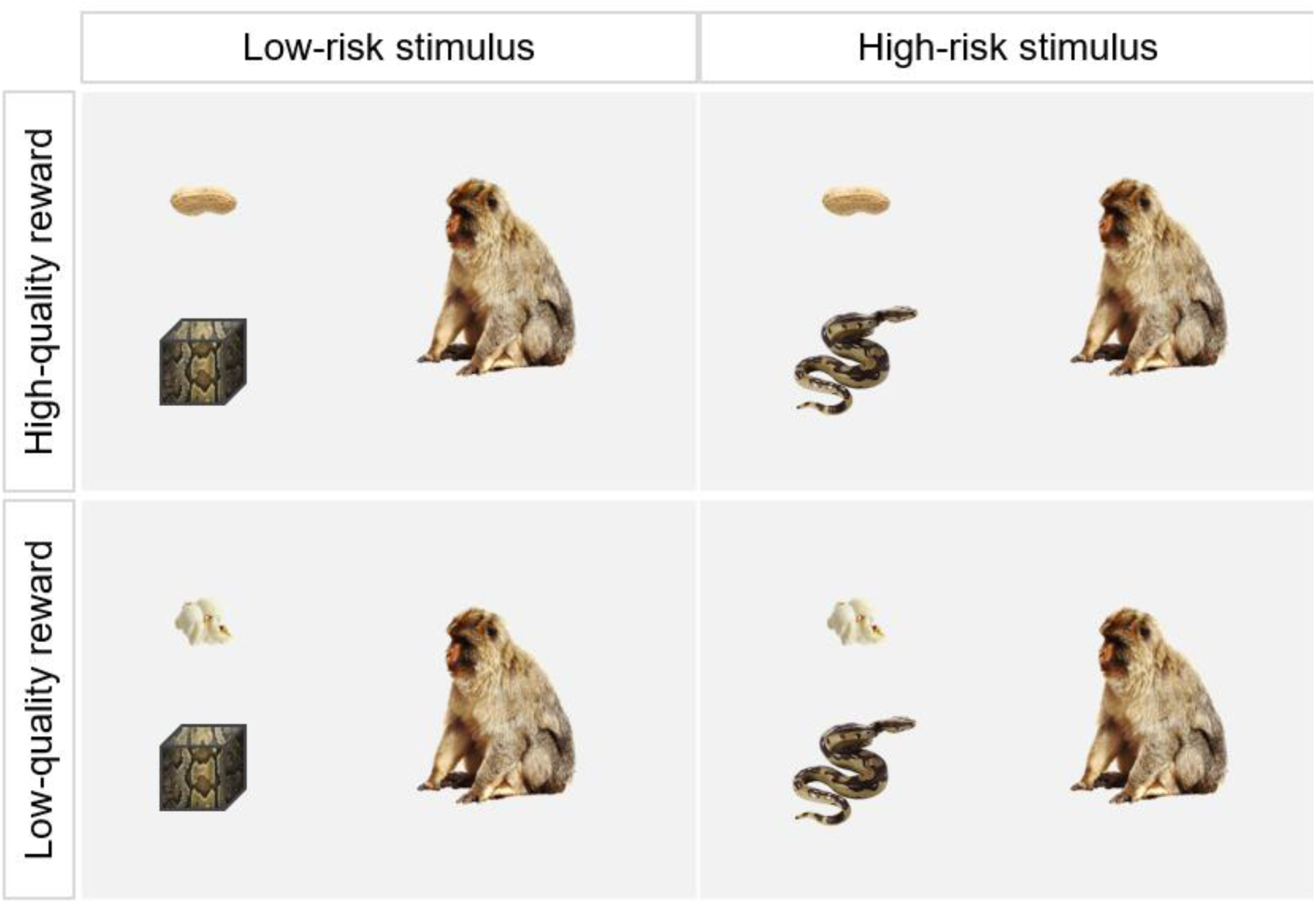
Schematic illustration of the four different combinations of high- and low-risk stimuli (snake and cube, respectively) and high- and low-quality rewards (peanut and popcorn, respectively).

### Familiarisation

Before the test trials, each individual went through a familiarisation phase, where only the food reward was placed in the wooden box, i.e., no risk stimulus, in order to rule out the possibility that the monkey was reluctant to approach the box itself and not the risk stimulus. The food reward that was presented to each subject depended on the food reward that the latter was attributed for the test trials. For example, if a subject was attributed snake and popcorn for the test trial, this subject would be presented with a popcorn during the familiarisation phase. Conversely, if a subject was attributed snake and peanut for the test trial, this subject would be presented with a peanut during the familiarisation phase.

Trials were conducted when no other monkeys were in close vicinity of the study subject. When the subject was located, the experimenter placed the wooden box on the floor. In order to elicit the subject’s interest in approaching the box and participate in the trial, the experimenter kneeled next to the box, extended her arm in the direction of the monkey, holding the attributed food reward. If the monkey approached, this was taken as a measure of motivation for the individual to participate in the task. Once the subject was within 1 to 2 meters of the experimental set-up, the experimenter slowly lowered the curtain, placed the food reward in the box, stood up and walked away. If the individual passed the familiarisation trial, i.e., retrieved the food reward, it could move on to the test trial.

### Test Trials

The procedure for the test trials was the same as in the familiarisation phase, but the wooden box would this time contain the risk stimulus in addition to the food reward. Similarly to the familiarisation phase, trials were conducted when the subject was relatively isolated from conspecifics, the attention of the subject was grabbed by showing them their attributed food reward, and the motion of the subject towards the experimental set-up was taken as an indicator of motivation for the individual to participate in the task. Once the subject was in closer proximity to the box, the experimenter lowered the curtain and placed the food reward next to the risk stimulus (at a distance of ∼30cm) which was attached to the box floor with wire or screws, for the snake and the cube, respectively. The monkey was then confronted with the decision of whether or not to retrieve the food reward placed next to the risk stimulus. Each monkey’s propensity to retrieve the food reward (yes or no) was taken as a measure of its risk-taking.

In the first study period, which took place between March and June of 2022, each individual monkey was assigned to one of the four possible factor combinations. In the second study period, from March to June 2023, each individual was assigned to the inverted combination. For instance, if a subject was assigned to snake and popcorn in 2022, the same individual would be assigned to cube and peanut in 2023. Food rewards and risk stimuli were balanced across sex and age categories. Data collection took place five to six days a week, and trials were always conducted early in the morning, before the first feeding. Across the two study periods, we conducted a total of 153 test trials, including 41 trials with cube and peanut, 35 trials with cube and popcorn, 41 trials with snake and peanut, and 36 trials with snake and popcorn. Taken together, 65 individuals participated in both study periods, 18 individuals participated in the first study period only, and five individuals participated in the second study period only.

### Data Coding

Each trial was filmed from two different angles. The first angle allowed to record the general scene of each trial, with a camera placed on a tripod and set up at a distance of 2 to 3 meters to the box. The second angle came from a camera attached behind the box, the lens of which fit through a hole cut out in the back panel of the box, allowing to record the monkeys’ frontal responses in the trials. Footage was recorded with GoPro HERO8 cameras. The videos were later coded in order to assess 1) whether the subject retrieved the reward, and 2) the latency of each subject to retrieve the food reward, taken as a measure of motivation to acquire the resource. Latency was measured from the moment the experimenter removed her hand from placing the food reward inside the box to the moment the monkey grabbed the reward. For the individuals that did not retrieve the food reward, the latency was recorded until the moment subjects walked away from the experimental box, turned their back to it, or demonstrated an apparent disinterest in it.

A second rater coded 20 of the videos in order to assess inter-rater reliability. We used Cohen’s kappa coefficient to assess inter-coder reliability in whether the subject retrieved the reward (*K* = 1, *N* = 20), and we used the inter-class correlation coefficient from the R package irr (version 0.84.1) to calculate inter-coder reliability in the latency of each subject to retrieve the food reward (*ICC* = 1, *N* = 20) (Koo & Li, 2016). Both reliability measurements indicate excellent reliability.

### Ethical Note

The study was exclusively non-invasive. The animals participated in the experiment on a voluntary basis, and the experimental protocol was approved by the management of La Forêt des Singes, Rocamadour, France. This research adhered to the ASAB/ABS Guidelines for the Use of Animals in Research (ASAB Ethical Committee/ABS Animal Care Committee, 2023).

### Statistical Analyses

To estimate how the probability of the animals to retrieve the reward was influenced by age, risk level, and reward quality, we fitted a Generalized Linear Mixed Model (GLMM; Baayen 2008) with binomial error structure and logit link function (McCullagh & Nelder 1989). We included age, risk level, and reward quality as well as all interactions among them up to the third order between them as fixed effects. We also included the fixed effects of sex and exposure (first or second exposure to the experiment) as control factors into the fixed effects part of the model. As we had two exposures for the majority of the individuals, we also included a random intercepts effect for the ID of the individual (random slopes could not be included because each individual was tested at most twice). Originally, we aimed to fit this model in a maximum likelihood framework but the respective model suffered from complete separation (Field 2005), convergence issues, and unrealistically large estimates (also for the random effect). Complete separation occurred as in one condition, namely the cube stimulus in combination with the peanut reward, the animals always retrieved the reward. We hence fitted the model in a Bayesian framework with regularising priors (McElreath 2020) (Model 1a). As priors for the fixed effects, we chose moderately uninformative priors, namely Gaussian priors with a mean of zero and a standard deviation of five. Such priors down-weight the importance of the data and make unrealistically large estimates less likely, yet still allow for the model to fit the observed response well. We further fitted three subset models of the full model with decreasing complexity in the fixed effects part each time removing interactions whose 95% credible interval encompassed the value of zero (Models 1b to 1d). We fitted these models using the function brm of the package brms (version 2.21.0; Bürkner, 2017, 2018, 2021) in R (version 4.3.3; R Core Team 2024). The sample analysed with these models comprised a total of 153 trials conducted with 88 individuals. Prior to fitting the model, we z-transformed age (Schielzeth, 2010) to ease model convergence.

In order to assess the factors influencing the animals’ latency to retrieve the reward, we fitted a model (Model 2a) with the same fixed and random effects structure as above-described Model 1. We excluded trials in which subjects did not retrieve the reward. Because of its skewed distribution, we log-transformed the response for this analysis. We fitted the model using a Gaussian error distribution and identity link function using the function lmer of the package lme4 (version 1.1-35.1; Bates et al. 2015). As an overall test of the effect of age and its interaction with risk level and reward quality, we compared the full model as just described with a null model lacking age, reward quality, risk level, and also all interactions among them. Such a full-null model comparison avoids ‘cryptic multiple testing’ (Forstmeier & Schielzeth 2011). We utilised a likelihood ratio test (Dobson 2002) for the full-null model comparison. To test the significance of individual fixed effects we used the Satterthwaite approximation to degrees of freedom as recommended by Luke (2017) as implemented in the function lmer of the package lmerTest (version 3.1-3; Kuznetsova et a. 2017). We estimated 95 % confidence intervals model estimates and fitted values y means of a parametric bootstrap (*N* = 1000 bootstraps; function bootMer of the package lme4). Inspection of a qq-plot of the residuals and residuals plotted against fitted values did not reveal strong deviations for the assumptions of normally distributed and homogeneous residuals. We determined model stability by dropping individuals, one at a time, fitting the full model to each of the subsets, and then comparing the range of estimates we obtained across the subsets with those the model on the full data set revealed. The same analysed with this model comprised 117 latencies obtained for 80 subjects. In case of Model 2a we fitted to subset models, one lacking the non-significant three-way interaction (Model 2b) and one lacking all non-significant interactions (Model 2c).

## RESULTS

In the first study period, we targeted 90 individuals and were able to test 83, as seven individuals did not approach the experimental set-up to participate in the task. In the second study period, we aimed at testing 80 individuals (10 subjects died from 2022 to 2023) and were able to test 70, as nine individuals did not approach the experimental set-up to participate in the task, and one individual’s trial was discarded due to interference by another monkey. In summary, we were able to run a total of 153 trials with 88 subjects (age range 3-31 years).

The probability to retrieve the reward was generally very high in all but one of the four combinations of stimulus and reward: individuals retrieved the reward in 117 out of 153 trials. More specifically, they took the peanut when paired with the cube in all of the 41 trials, and in 34/41 trials when paired with the snake. They took the popcorn in 30/35 cases when paired with the cube, but only in 12/36 cases when paired with the snake (Figure 2).

**Figure 2.**
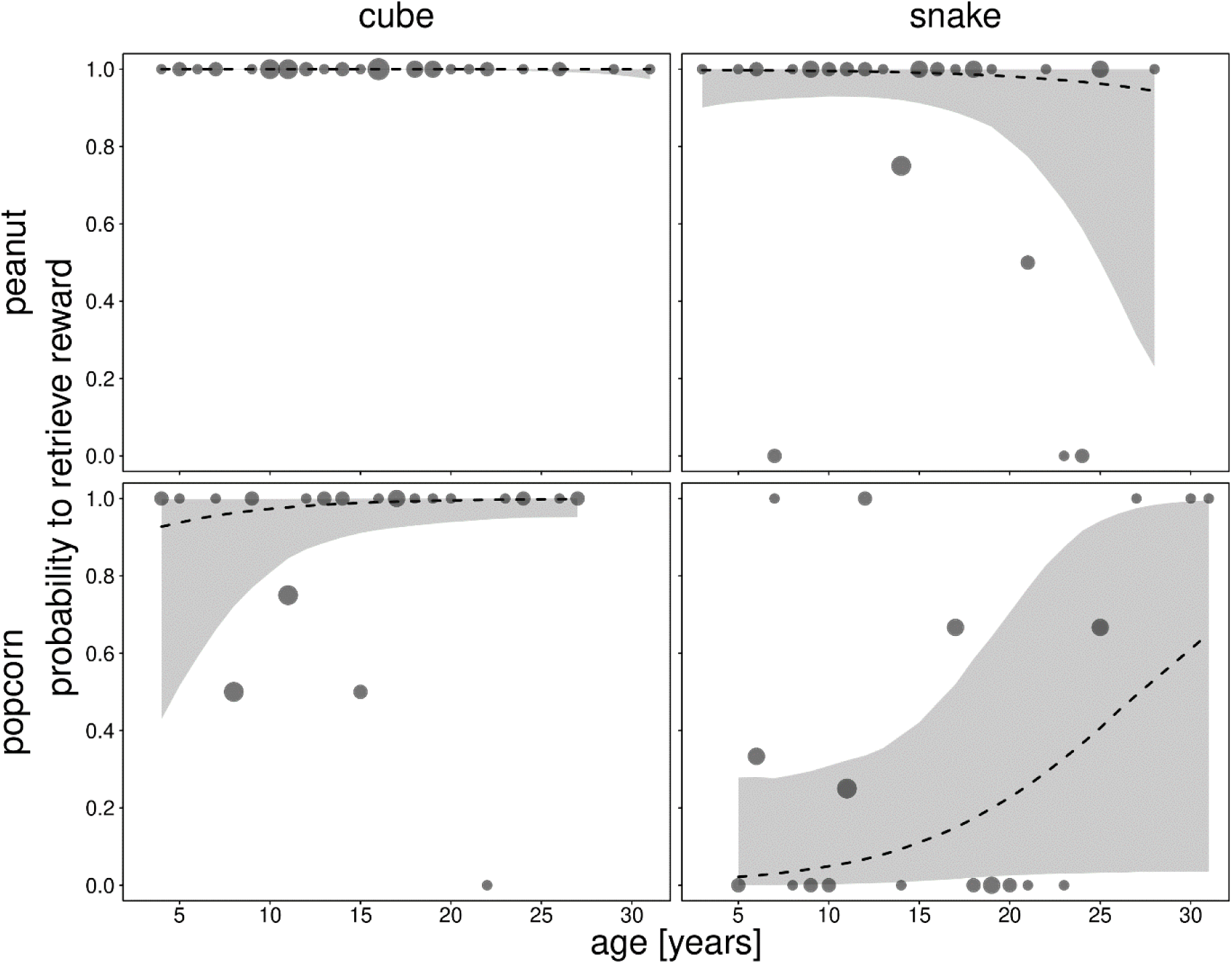
Probability to retrieve the reward as a function of age and separately for different stimuli (columns) and rewards (rows). Dots show the average probabilities per age whereby their area is proportionate to the number of trials per combination of age, stimulus, and reward (range 1 to 5 trials). The dashed lines with grey polygons depict the fitted model and its 95% credible interval for all other terms being centred (Model 1a).

The results of the Bayesian models showed no evidence for an effect of the three-way interaction between age, reward, and stimulus (Model 1a, 95% credible interval [-3.098, 7.357]). We therefore removed the three-way interaction and fitted the model again with three two-way interactions (Model 1b, reward*age + stimulus*age + reward*stimulus). The results of this model showed no evidence for an effect of the interactions between stimulus and age (95% credible interval [-2.843, 2.939]), and between stimulus and reward (95% credible interval [-10.111, 2.359]). However, there was weak evidence for an effect of the interaction between reward and age (95% credible interval [-0.002, 5.890]) (see full results for Models 1a and 1b in Supplementary Tables 1 and 2, respectively). Therefore, we fitted another model excluding the interactions between stimulus and age, and stimulus and reward, but retaining the interaction between reward and age (Model 1c). This model showed evidence for an effect of stimulus, whereby subjects were less likely to retrieve the reward when paired with the snake rather than with the cube (95% credible interval [-10.889, -3.618]). There was also weak and not very conclusive evidence for an effect of the interaction between reward and age (95% credible interval [-0.070, 4.932]), whereby the probability for the individuals to retrieve a peanut did not vary with age, but the probability to retrieve a popcorn increased with age. However, the uncertainty of the slope estimate was very high (Table 1; Figure 2). Nonetheless, these results go along the lines of the significant interaction of reward and age observed in the latency analysis (see below), thus providing support for this model (i.e., Model 1c). Finally, we refitted the model excluding the interaction between reward and age (Model 1d), and this model provided clear evidence for an effect of reward (95% credible interval [-11.269, -3.295]), and stimulus (95% credible interval [-10.324, - 3.312]), whereby subjects were less likely to retrieve the reward when it was a popcorn rather than a peanut, and less likely to retrieve the reward when paired with the snake rather than the cube (Table 2).

**Table 1.** Results of the reduced Bayesian model with the animals’ probability of retrieving the reward as the response, retaining only the interaction between age and reward quality (Model 1c).

| Term | Estimate | SE | CIlower | Clupper | Rhat | BulkESS | TailESS |
| --- | --- | --- | --- | --- | --- | --- | --- |
| Intercept | 9.947 | 2.881 | 5.336 | 16.559 | 1.002 | 3094.856 | 3694.944 |
| Reward | -6.841 | 1.997 | -11.268 | -3.550 | 1.002 | 3125.340 | 3949.746 |
| Age | -0.788 | 1.055 | -2.972 | 1.193 | 1.001 | 5709.561 | 4297.833 |
| Stimulus | -6.691 | 1.866 | -10.889 | -3.618 | 1.002 | 3513.213 | 4481.191 |
| Sex | 1.319 | 1.421 | -1.411 | 4.272 | 1.001 | 6349.267 | 5206.582 |
| Exposure | 1.619 | 1.131 | -0.418 | 4.066 | 1.001 | 6717.183 | 5028.132 |
| Reward:Age | 2.122 | 1.250 | -0.070 | 4.932 | 1.001 | 5276.808 | 4739.667 |

Indicated are posterior means, together with their standard errors and 95% credible intervals. Age was z-transformed to a mean of zero and a standard deviation (sd) of one; mean and sd of the original age were 15.0 and 6.7 years. All factors were dummy coded; reference levels were female (sex), 1 (exposure), peanut (reward), and cube (stimulus).

**Table 2.** Results of the reduced Bayesian model with the animals’ probability of retrieving the reward as the response, lacking all interactions (Model 1d).

| Term | Estimate | SE | CIlower | Clupper | Rhat | BulkESS | TailESS |
| --- | --- | --- | --- | --- | --- | --- | --- |
| Intercept | 9.340 | 2.816 | 4.943 | 16.012 | 1.001 | 2170.930 | 2900.818 |
| Reward | -6.556 | 2.009 | -11.269 | -3.295 | 1.001 | 2253.977 | 2949.115 |
| Age | 0.647 | 0.681 | -0.605 | 2.088 | 1.001 | 3896.567 | 3803.154 |
| Stimulus | -6.193 | 1.771 | -10.324 | -3.312 | 1.001 | 2530.167 | 3054.499 |
| Sex | 1.235 | 1.400 | -1.559 | 4.149 | 1.001 | 4262.427 | 3762.435 |
| Exposure | 1.557 | 1.087 | -0.336 | 3.891 | 1.001 | 5191.080 | 4273.857 |

Indicated are posterior means, together with their standard errors and 95% credible intervals.
Age was z-transformed to a mean of zero and a standard deviation (sd) of one; mean and sd of the original age were 15.0 and 6.7 years. All factors were dummy coded; reference levels were female (sex), 1 (exposure), peanut (reward), and cube (stimulus).

To gain further insights into the animals’ decision-making process, we analysed the latency to take the reward for the 117 trials where the animals did so (*N* = 80 subjects, age range 3-31 years). The full-null model comparison indicated a significant effect of the predictor variables (full-null model comparison Model 2a, *χ*^2^ = 36.343, *df* = 7, *P* < 0.001). The model did not reveal a significant effect of the three-way interaction, however (*P* = 0.374); we hence removed this term and proceeded with including the two-way interactions (Model 2b). We found a significant interaction between reward and age (*P* = 0.013), but no significant interaction between stimulus and age (*P* = 0.13) (see full results for Models 2a and 2b in Supplementary Tables 3 and 4, respectively). We therefore fitted a final model with only the two-way interaction between reward and age and the main factors (Model 2c, Table 3). The interaction between reward and age was still significant (*P* = 0.026). Plotting the data and the model revealed no effect of age on the latency to take the peanut, but younger individuals took more time to take the popcorn than older animals, contrary to our predictions (Figure 3). We also observed a significant main effect of the stimulus on the latency to take the reward (*P* < 0.001), with animals taking more time when confronted with the snake (median = 7.1 s) than with the cube (median = 4.5 s). Finally, males were faster to take the reward (median = 4.8 s) than females (median = 6.1 s, *P* = 0.029).

**Figure 3.**
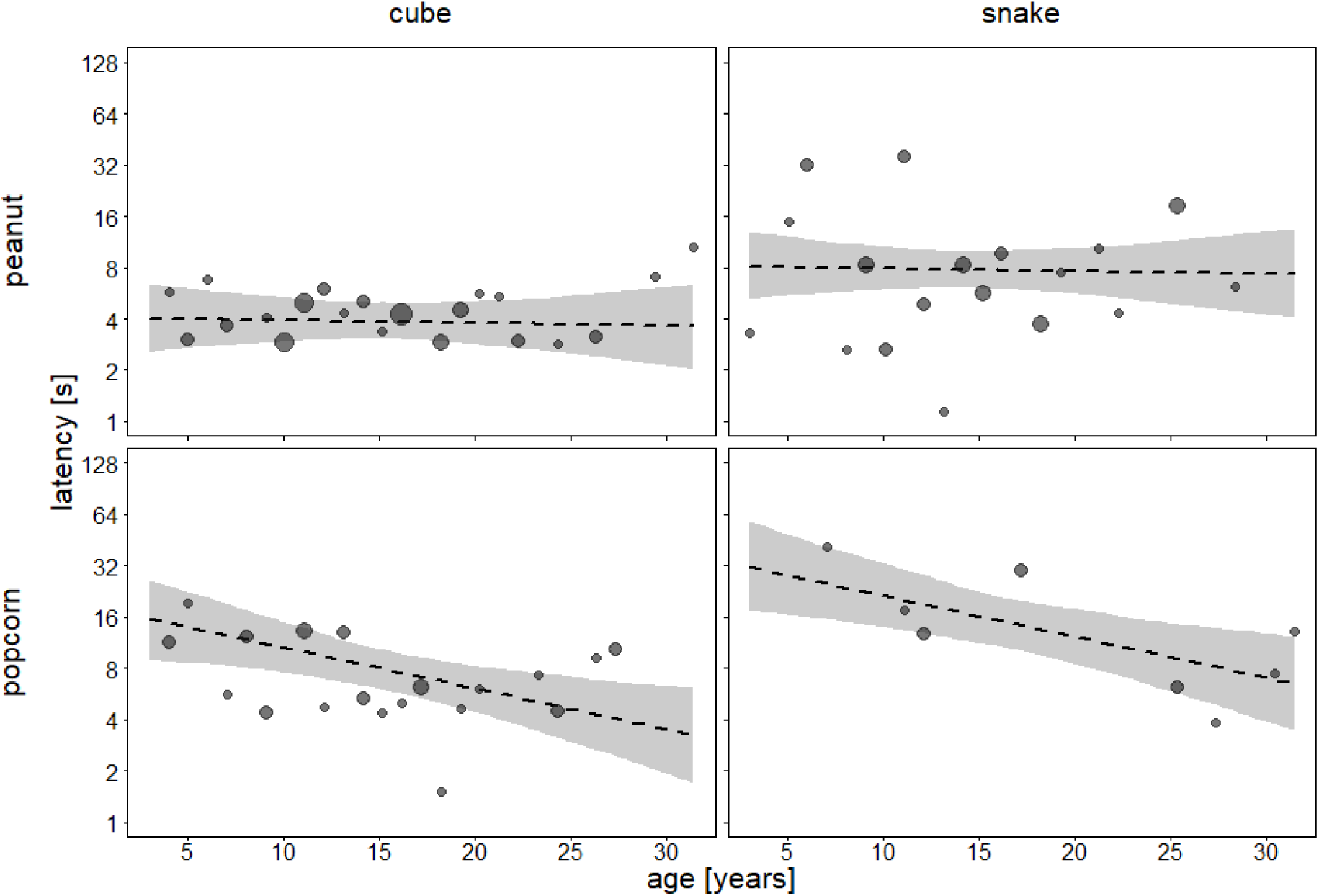
Latency to retrieve the reward as a function of age for the different stimuli and rewards. Dots show the averaged latencies per age whereby their area is proportionate to the number of trials per combination of age, stimulus, and reward (range 1 to 5 trials). The dashed lines with grey polygons depict the fitted model and its 95% credible interval for all other terms being centred (Model 2c). Note the logarithmic scale of the y-axis.

**Table 3.**
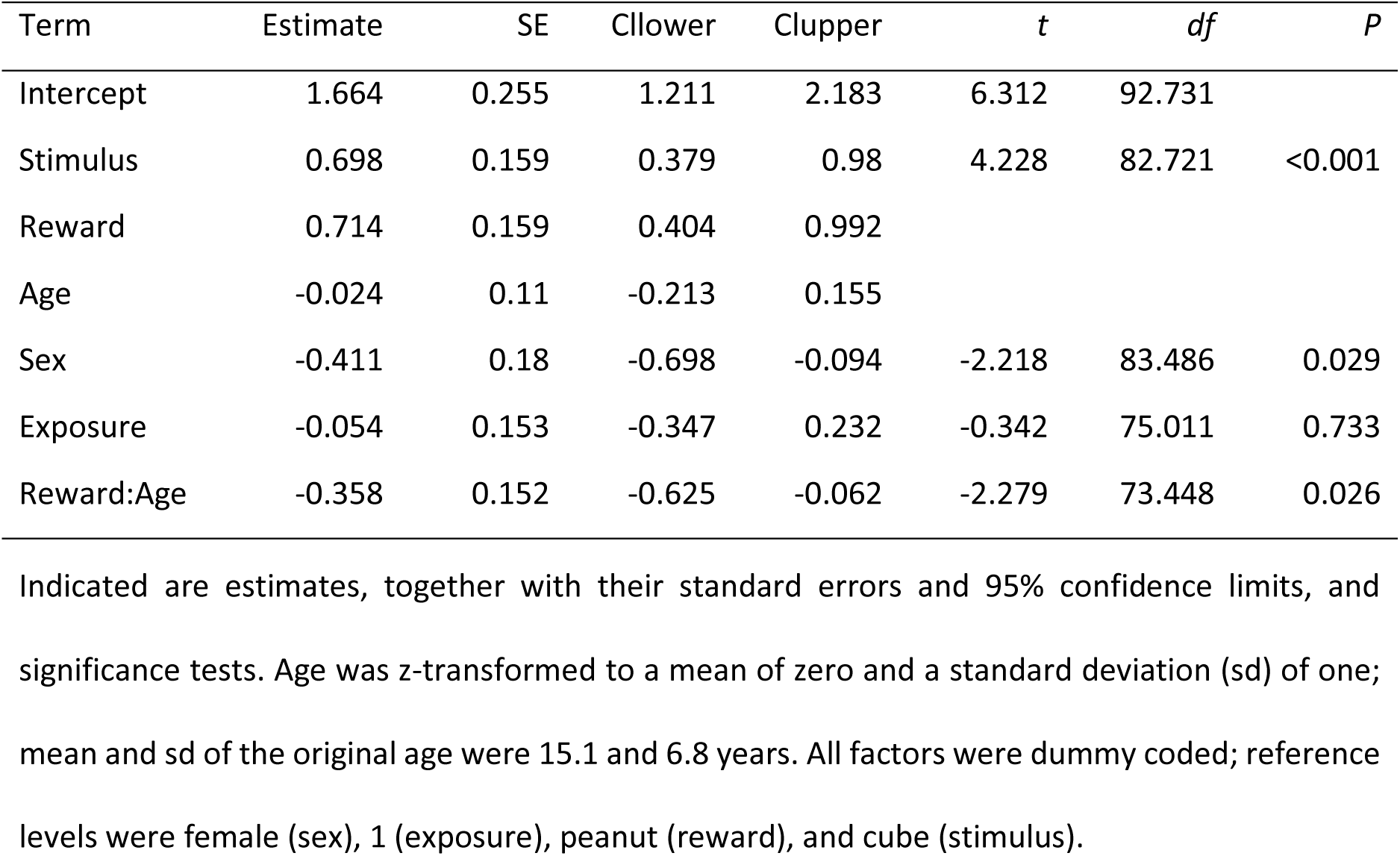
Results of the reduced model with the animals’ latency to retrieve the reward as a response, retaining only the interaction between age and reward quality (Model 2c).

| Term | Estimate | SE | CI lower | CI upper | *t* | *df* | *P* |
| --- | --- | --- | --- | --- | --- | --- | --- |
| Intercept | 1.664 | 0.255 | 1.211 | 2.183 | 6.312 | 92.731 |  |
| Stimulus | 0.698 | 0.159 | 0.379 | 0.98 | 4.228 | 82.721 | <0.001 |
| Reward | 0.714 | 0.159 | 0.404 | 0.992 |  |  |  |
| Age | -0.024 | 0.11 | -0.213 | 0.155 |  |  |  |
| Sex | -0.411 | 0.18 | -0.698 | -0.094 | -2.218 | 83.486 | 0.029 |
| Exposure | -0.054 | 0.153 | -0.347 | 0.232 | -0.342 | 75.011 | 0.733 |
| Reward:Age | -0.358 | 0.152 | -0.625 | -0.062 | -2.279 | 73.448 | 0.026 |

Indicated are estimates, together with their standard errors and 95% confidence limits, and significance tests. Age was z-transformed to a mean of zero and a standard deviation (sd) of one; mean and sd of the original age were 15.1 and 6.8 years. All factors were dummy coded; reference levels were female (sex), 1 (exposure), peanut (reward), and cube (stimulus).

## DISCUSSION

We found weak and not very conclusive evidence supporting age-related variation in ecological risk-taking when considering the probability to take the reward. Due to issues with complete separation of the data and ceiling effects in the analysis of the probability to take the reward, the results of the statistical analysis need to be treated with a pinch of salt, however. The model results suggest that regardless of age, there was an effect of the main factors risk level and reward quality on the monkeys’ propensity to retrieve the reward. Individuals were overall more likely to take the peanut than the popcorn and less likely to take the reward when paired with the snake than the cube. While our results hint at a higher inclination for older individuals to retrieve the reward when popcorn was paired with snake, than their younger conspecifics, the credible intervals of fitted values were very wide. Yet, we found no support for our working hypothesis that older animals would be less risk prone than younger ones.

The analysis of the latency to retrieve the reward was more informative and clearly showed that the time to decision was affected by the stimulus and the interaction between reward and age. As predicted, animals were more cautious when confronted with the snake than with the cube. Contrary to our predictions, however, older individuals were faster than their younger conspecifics to retrieve the reward when it was popcorn, even when paired with the snake. This finding contrasts with the previous observation that older Barbary macaques showed higher neophobia with older age (Almeling et al., 2016), but may be explained by the presence of the food reward (see also Almeling et al. 2016).

One point to consider is that the animals did not always appear to have detected the snake before they took the reward (e.g., see Video 5). In these cases, we cannot be sure whether the subject would have retrieved the reward had it seen the risky object first. While these instances were few and had no influence on our overall results, they did present an unexpected caveat in the proceedings of our study.

The fact that older subjects were faster to take the reward may be explained by the fact that animals aged 20 and older are typically lower ranking (Rathke & Fischer, 2021). Consequently, they have more limited access to resources than their younger conspecifics (Sterck & Steenbeek, 1997). Under normal circumstances, foods are regularly usurped or monopolised by younger and more dominant individuals, preventing old monkeys to profit from them. The conditions provided in our experiment, namely relative social isolation of the subject and an “open” access to a food reward, may have enticed those older, lower-ranking, individuals to reach for an otherwise inaccessible resource. Against the food competition among these animals, older individuals may have been more willing to take a risk, even for a lower-quality reward, for the potentially scarce opportunity to acquire an infrequently occurring resource. Moreover, and in contrast to the experiments done by Almeling and colleagues (2016), the access of the food reward did not involve problem-solving, and only subjects that had passed the familiarisation test participated in the trials, thus excluding the effects of neophobia. The high prevalence of taking the food when paired with the cube also indicates that the snake pattern we chose constituted only a mildly aversive stimulus, at best. However, rank-associated access to food is not the only explanation for the shorter latencies in old age, as young individuals are normally lower ranking than middle-aged ones, and these individuals showed some of the longest latencies to take the reward. Thus, in conclusion, a combination of risk aversion, experience, and an increased value of a food resource in old age may best explain the observed pattern.

Our results complement previous findings showing that primates take into account the social context of a given situation and are able to alter their risk-taking strategies accordingly (Haux et al., 2021). For instance, tufted capuchins are more prone to taking risks when alone than in the presence of a conspecific (Zoratto et al., 2018). Considering that the costs of a physical altercation could be higher for older individuals whose body condition might be more fragile (Nussey et al., 2013), older monkeys may have had a higher incentive to take risks when alone than younger ones. Our results therefore suggest that the value of a resource is not fixed, it is possible that the value that is attributed to a certain resource, and the motivation to acquire it, changes with age, but not always in the predicted way.

Overall, our findings are in keeping with most previous studies investigating age-related changes in risk-taking in primates, although sample sizes and age ranges are typically small, precluding strong inferences (reviewed in De Petrillo & Rosati, 2021). The design of our study, however, contrasts with many previous studies on decision-making under risk in primates, which have focused on methodologies that involve the animals to be trained in order to partake in the desired tasks. Animals often have to learn various payoffs associated with ‘safe’ and ‘risky’ options (e.g., Proctor et al., 2014), or learn to exchange tokens or food items with human experimenters (e.g., Broihanne et al., 2019). However, for foraging animals, potential risk manifests not necessarily in short-term variation of statistical patterns frequently used in studies framed within economic theories. Instead, animals must decide whether to take a certain route, although lions have been seen in that area earlier, or they have to consider whether or not they wade through water bodies possibly infested with crocodiles (Cheney & Seyfarth, 2008). As such, and in line with the notion that the ecological validity of a task may influence the expression of risk preferences (Eisenreich et al., 2019), we investigated risk attitudes using a more naturalistic “one-shot” approach.

While subjects in the more ‘economically oriented’ studies typically receive a certain amount of reward, even in the risky options, natural conditions include the possibility of receiving nothing at all. In this sense, the naturalistic approach to risk in our study is two-fold: firstly, by the snake that evokes a potential predatory hazard; secondly, by the fact that subjects may walk away from the task empty-handed as a result of a missed opportunity to acquire the offered resource. In view of this concept, the animals must decide which risk to take or avoid, and our experiment illustrates the assessment of this trade-off. Our results also further corroborate the notion that the nature of the reward appears to impact risk attitudes (Heilbronner & Hayden, 2013). In humans too, people’s risk preferences vary across decisions involving food, prizes, or money, based on the perceived value and usefulness of the rewards (Rosati & Hare, 2016). By considering the value of potential losses and gains, as well as the biological relevance of the context that a given risk presents itself in (Babb et al., 2024; Haux et al., 2021), we may be better able to determine the parameters guiding primate risk-taking strategies in the ecological domain.

In conclusion, our study does not support the idea that risk-taking propensity invariably declines with age in primates. Instead, the potential costs and benefits of a given decision may vary with the type of decision at hand. Overall, Barbary macaques across age seem to assess these cost-benefit trade-offs in their decisions involving risk in the ecological domain.

To get a better grasp on the expression of risk in primates, follow-up research could present a wider variation of reward and risk items, as a way to pinpoint how the salience of the stimuli may guide individuals’ threshold for risk propensity. Would the animals still reach for the peanut if paired with a snake that was larger, or one that was moving? Would they be more willing to approach the rubber snake if the popcorn came in a handful and not a single piece? A further step could be to introduce a certain level of ambiguity by providing the animals with less information regarding the reward and risk stimuli as foraging with incomplete knowledge may better reflect situations they face under natural conditions. In addition, future studies could bear a greater focus on the latency of animals to engage in a risky choice since – as suggested with our results – the latter may hold further detailed information about the subjects’ deliberation and decision-making processes. Delving deeper into such observations may tell us more about how monkeys assess the value of gains against potential adverse outcomes in risky foraging events.

## Author Contributions

**Tiffany C. Bosshard**: Conceptualization, Methodology, Investigation, Data curation, Writing (original draft). **Julia Fischer**: Conceptualization, Methodology, Writing (review and editing). **Roger Mundry**: Formal analysis, Writing (review and editing).

## Data availability

The data and statistical code associated with this article can be found at https://osf.io/vjmbr/?view_only=7d20676f9f8a4fb88a5915ceb3662872.

## Acknowledgments

We thank Ellen Merz, Gilbert de Turckheim, and Roland Hilgartner for their permission to carry out this study at La Forêt des Singes. We also thank the dedicated staff members of the park for their invaluable assistance and support throughout this project. We are particularly grateful to Victor Gass, Nadja Vögtle, and Claire des Pallières for their assistance in collecting the data in the field. We also thank Dr. Derek Murphy for his invaluable support throughout the statistical analyses. This research was funded by the Deutsche Forschungsgemeinschaft (DFG, German Research Foundation) – Project-ID 254142454/GRK 2070 “Understanding Social Relationships”.

## Supplementary Material

Supplementary material associated with this article can be found in the online version.

## Supplementary material

Bosshard, Mundry, Fischer. Ecological risk-taking across age in Barbary macaques.

**Supplementary Table 1.** Results of the full model, with the animals’ probability of retrieving the reward as the response, with the three-way interaction of age, reward, and stimulus (Model 1a).

| Term | Estimate | SE | CIlower | CIupper | Rhat | BulkESS | TailESS |
| --- | --- | --- | --- | --- | --- | --- | --- |
| Intercept | 10.806 | 3.144 | 5.407 | 17.673 | 1.002 | 2901.701 | 4307.674 |
| Reward | -6.397 | 2.406 | -11.579 | -2.072 | 1.000 | 3876.995 | 4307.553 |
| Stimulus | -5.982 | 2.298 | -10.771 | -1.828 | 1.000 | 3855.601 | 4879.926 |
| Age | -0.328 | 1.988 | -4.307 | 3.62 | 1.002 | 3206.412 | 4145.536 |
| Sex | 1.651 | 1.792 | -1.742 | 5.347 | 1.001 | 3482.328 | 4406.424 |
| Exposure | 2.095 | 1.414 | -0.295 | 5.233 | 1.001 | 5240.562 | 4535.698 |
| Reward:Stimulus | -3.525 | 3.282 | -10.434 | 2.36 | 1.000 | 3564.971 | 4667.915 |
| Reward:Age | 1.479 | 2.187 | -2.742 | 5.977 | 1.002 | 3119.502 | 4669.562 |
| Stimulus:Age | -1.023 | 2.093 | -5.254 | 2.959 | 1.001 | 3345.661 | 4299.735 |
| Reward:Stimulus:Age | 2.038 | 2.665 | -3.098 | 7.357 | 1.001 | 3354.458 | 4633.341 |

Indicated are posterior means, together with their standard errors and 95% credible intervals. Age was z-transformed to a mean of zero and a standard deviation (sd) of one; mean and sd of the original age were 15.0 and 6.7 years. All factors were dummy coded; reference levels were female (sex), 1 (exposure), peanut (reward), and cube (stimulus).

**Supplementary Table 2.** Results of the reduced model with the animals’ probability of retrieving the reward as the response, lacking the three-way interaction of predictor variables (Model 1b).

| Term | Estimate | SE | CIlower | CIupper | Rhat | BulkESS | TailESS |
| --- | --- | --- | --- | --- | --- | --- | --- |
| Intercept | 10.574 | 3.195 | 5.286 | 17.576 | 1.001 | 3486.1 | 4338.311 |
| Reward | -6.226 | 2.419 | -11.315 | -1.92 | 1.001 | 4551.621 | 4330.89 |
| Stimulus | -5.969 | 2.28 | -10.813 | -1.899 | 1.002 | 4729.88 | 4987.407 |
| Age | -0.999 | 1.754 | -4.459 | 2.49 | 1.000 | 3675.046 | 3978.784 |
| Sex | 1.677 | 1.745 | -1.683 | 5.441 | 1.001 | 4940.41 | 4947.302 |
| Exposure | 2.006 | 1.408 | -0.395 | 5.111 | 1.000 | 6436.059 | 4307.225 |
| Reward:Stimulus | -3.282 | 3.182 | -10.111 | 2.359 | 1.001 | 4222.928 | 5572.239 |
| Reward:Age | 2.653 | 1.497 | -0.002 | 5.89 | 1.000 | 3923.266 | 4071.138 |
| Stimulus:Age | 0.153 | 1.45 | -2.843 | 2.939 | 1.001 | 4772.354 | 4932.774 |

Indicated are posterior means, together with their standard errors and 95% credible intervals. Age was z-transformed to a mean of zero and a standard deviation (sd) of one; mean and sd of the original age were 15.0 and 6.7 years. All factors were dummy coded; reference levels were female (sex), 1 (exposure), peanut (reward), and cube (stimulus).

**Supplementary Table 3.** Results of the full model, with the animals’ latency to retrieve the reward as a response, with the three-way interaction of age, reward, and stimulus (Model 2a).

| Term | Estimate | SE | Clower | Clupper | *t* | *df* | *P* | min | max |
| --- | --- | --- | --- | --- | --- | --- | --- | --- | --- |
| Intercept | 1.744 | 0.258 | 1.271 | 2.229 | 6.456 | 96.766 |  | 1.665 | 1.819 |
| Reward | 0.551 | 0.203 | 0.166 | 0.893 |  |  |  | 0.466 | 0.593 |
| Stimulus | 0.531 | 0.199 | 0.17 | 0.888 |  |  |  | 0.431 | 0.611 |
| Age | 0.012 | 0.138 | -0.252 | 0.25 |  |  |  | -0.035 | 0.047 |
| Sex | -0.434 | 0.177 | -0.751 | -0.148 | -2.346 | 81.56 | 0.021 | -0.483 | -0.358 |
| Exposure | -0.05 | 0.15 | -0.337 | 0.227 | -0.318 | 73.768 | 0.751 | -0.094 | 0.019 |
| Reward:Stimulus | 0.611 | 0.379 | -0.009 | 1.26 | 1.542 | 89.561 | 0.127 | 0.378 | 0.784 |
| Reward:Age | -0.261 | 0.209 | -0.62 | 0.091 | -1.194 | 106.371 | 0.235 | -0.318 | -0.174 |
| Stimulus:Age | -0.101 | 0.213 | -0.483 | 0.277 | -0.452 | 106.576 | 0.652 | -0.271 | 0.036 |
| Reward:Stimulus:Age | -0.325 | 0.349 | -0.894 | 0.288 | -0.893 | 80.62 | 0.374 | -0.455 | -0.152 |

Indicated are estimates, together with their standard errors and 95% confidence limits, and significance tests. Age was z-transformed to a mean of zero and a standard deviation (sd) of one; mean and sd of the original age were 15.1 and 6.8 years. All factors were dummy coded; reference levels were female (sex), 1 (exposure), peanut (reward), and cube (stimulus).

**Supplementary Table 4.** Results of the reduced model, with the animals’ latency to retrieve the reward as a response, lacking the three-way interaction of predictor variables (Model 2b).

| Term | Estimate | SE | Clower | Clupper | *t* | *df* | *P* |
| --- | --- | --- | --- | --- | --- | --- | --- |
| Intercept | 1.738 | 0.259 | 1.255 | 2.224 | 6.441 | 97.331 |  |
| Reward | 0.546 | 0.204 | 0.187 | 0.89 |  |  |  |
| Stimulus | 0.52 | 0.199 | 0.176 | 0.848 |  |  |  |
| Age | 0.069 | 0.124 | -0.166 | 0.27 |  |  |  |
| Sex | -0.439 | 0.178 | -0.728 | -0.153 | -2.376 | 82.738 | 0.02 |
| Exposure | -0.042 | 0.15 | -0.339 | 0.263 | -0.273 | 73.83 | 0.786 |
| Reward:Stimulus | 0.582 | 0.38 | -0.013 | 1.184 | 1.472 | 89.479 | 0.145 |
| Reward:Age | -0.398 | 0.15 | -0.685 | -0.114 | -2.535 | 73.532 | 0.013 |
| Stimulus:Age | -0.242 | 0.151 | -0.526 | 0.055 | -1.533 | 73.275 | 0.13 |

Indicated are estimates, together with their standard errors and 95% confidence limits, and significance tests. Age was z-transformed to a mean of zero and a standard deviation (sd) of one; mean and sd of the original age were 15.1 and 6.8 years. All factors were dummy coded; reference levels were female (sex), 1 (exposure), peanut (reward), and cube (stimulus).

